# Biomarkers of lesion severity in a rodent model of nonarteritic anterior ischemic optic neuropathy (rNAION)

**DOI:** 10.1101/2020.11.18.388132

**Authors:** Yan Guo, Zara Mehrabian, Mary A. Johnson, Neil R. Miller, Amanda D. Henderson, John Hamlyn, Steven L. Bernstein

## Abstract

The rodent model of nonarteritic anterior ischemic optic neuropathy (rNAION) is similar in many of its pathophysiological responses to clinical NAION. However, little is known of the parameters associated with rNAION induction severity and if pre- or early post-induction biomarkers can be identified that enable prediction of lesion severity and ultimate loss of retinal ganglion cells (RGCs). Adult male Sprague-Dawley outbred rats were evaluated for various parameters including physiological characteristics (heart rate, respiratory rate, temperature, hematocrit), optic nerve head (ONH) appearance, pre- and post-induction mean diameter, and intravenous fluorescein and indocyanine green angiographic patterns of vascular leakage at 5 hours post-induction, performed using a spectral domain-optical coherence tomography (SD-OCT) instrument. These parameters were correlated with ultimate RGC loss by Brn3a (+) immunohistology. RGC loss also was correlated with the relative level of laser exposure. The severity of ONH edema 2d, but not 5hr, post induction was most closely associated with the degree of RGC loss, revealing a threshold effect, and consistent with a compartment syndrome where a minimum level of capillary compression within a tight space is responsible for damage. RGC loss increased dramatically as the degree of laser exposure increased. Neither physiological parameters nor the degree of capillary leakage 5hr post induction were informative as to the ultimate degree of RGC loss. Similar to human NAION, the rNAION model exhibits marked variability in lesion severity. Unlike clinical NAION, pre-induction ONH diameter likely does not contribute to ultimate lesion severity; however, cross-sectional ONH edema can be used as a biomarker 1-2d post-induction to determine randomization of subjects prior to inclusion in specific neuroprotection or neuroregeneration studies.

## Introduction

Nonarteritic anterior ischemic optic neuropathy (NAION) is the most common cause of sudden optic nerve-related vision loss in individuals over the age of 50 in the developed world. In this condition, vascular decompensation occurring in the tightly restricted space of the optic nerve head (ONH) results in a compartment syndrome and sudden ischemia in the anterior portion of the ON. Human NAION is distinguished by ONH capillary incompetence, demonstrable by both intravenous fluorescein angiography (IVFA) and indocyanine green (ICG) leakage in early stages of the disease [1]. Direct tissue analysis of this disease is difficult to obtain, as NAION is a nonfatal disease. Clinical diagnosis instead relies on symptoms, signs, and electrophysiological findings (eg, visual evoked potentials (VEP) [2, 3].

To circumvent the problems inherent in NAION clinical research, we developed rodent and primate nonarteritic anterior ischemic optic neuropathy (rNAION and pNAION, respectively) [4–6]. These models are pathophysiologically similar in many ways to the clinical disease, in terms of presentation with ONH edema, acellular and cellular inflammatory responses, ON-axon loss, isolated retinal ganglion cell (RGC) loss and visual debilitation [7]. In these models, laser-induced superoxide radicals from a photosensitive dye induce anterior ON capillary decompensation and leakage, ultimately resulting in ON edema, a compartment syndrome, and a lesion resembling the clinical one [8]. We previously documented fluorescein leakage in rNAION [5].

Despite extensive research using murine models of NAION [4, 9, 10], many questions still remain concerning rNAION induction consistency, predictability of lesion severity, induction parameters, and if different NAION biomarkers, such as post-induction ON edema or capillary leakage, correlate with ultimate RGC and vision loss [11]. This problem is homologous to the variability in clinical lesion severity seen in NAION as well as in other central nervous system (CNS) ischemia/stroke models [12, 13]. This variability results in the statistical requirement to analyze large numbers of test animals to determine efficacy of proposed neuroprotective agents, thus requiring considerable time, effort, and cost. Identification of relevant early NAION biomarkers might improve selection of equivalently affected test animals prior to treatment randomization, potentially improving predictive accuracy of neuroprotective treatments, while using fewer animals.

In an attempt to improve lesion consistency in our rNAION model, we evaluated multiple variables potentially associated with lesion severity in an outbred strain with attention paid to the response variations between animals. Variables included correlation of pre-induction rodent ONH diameter with post-induction edema, because the risk of clinical NAION is associated with a small ONH (the ‘disk at risk’), as well as the intrinsic variables of heart rate and body temperature at induction. Additionally, we evaluated physical parameters of the ONH early (5hr) and late (2d) post-induction as potential biomarkers of model severity. These included: 1) ONH appearance and degree of ONH edema by direct visualization and spectral domain-optical coherence tomography (SD-OCT) and 2) vascular decompensation and capillary leakage.

As the rNAION lesion largely results from a compartment syndrome following vascular decompensation and leakage [3, 14], we reasoned that evaluation of early (3-5hr) vascular leakage signal intensity could be used as a potential biomarker of lesion severity. Five hours post induction was chosen because we previously demonstrated that a neuroprotective rNAION treatment (PGJ_2_) started at this time point was effective [14]. Leakage analysis used two different dyes (fluorescein and indocyanine green) because of their different properties. Because fluorescein is excited by visible light (peak 494nm), evaluation of deep/pigmented structures is limited. Additionally fluorescein binds poorly to serum proteins [15], diffusing away from damage sites. In contrast, ICG is excited by infrared light (peak ~780nm), enabling deeper tissue penetration and imaging through pigmented tissue (16). ICG also is ~98% bound to serum lipoproteins [17], slowing diffusion through tissues and enabling identification of focal subzones of leakage. The signals from both dyes can be analyzed using the widely available Heidelberg SD-OCT device (Heidelberg Engineering, Franklin, MA). We correlated early potential biomarkers with later (2d) ONH edema and ultimate (30d) RGC loss.

Sodium-potassium pumps (Na pumps) play the critical role in the maintenance of monovalent ion gradients across the plasma membrane. Under normal conditions, Na pumps hydrolyze ~50-55% of the brain adenosine triphosphate (ATP) [18, 19]. During early ischemia, declining cellular ATP results in reduced Na pump activity, diminished ion gradients, and subsequent edema formation [20]. Agents minimizing the loss of Na pump activity thus would be expected to ameliorate the adverse impact of ischemia and inhibit edema. Endogenous cardiotonic steroids (CTS) such as ouabain and digoxin-like compounds that inhibit Na pumps have been described in the CNS [21, 22]. During ischemia, CTS may leak from anoxic cells and accumulate in the extracellular space in concentrations that inhibit Na pumps in the local environment, resulting in extracellular fluid leakage and edema. In the final part of this study, we evaluated if systemically blocking endogenous CTS using the commercially available anti-ouabain antibody (DigiFab) would be effective in reducing ONH edema and leakage.

## Methods

### Animals

All animal protocols were approved by the UMB institutional animal care and use committee. Male Sprague-Dawley rats (200-250g) were used in this study.

### Anesthesia and rNAION induction

Animals were anesthetized with an intraperitoneal mixture of ketamine and xylazine (80mg; 4mg/kg) and kept on warming pads during anesthesia induction, rNAION induction, and recovery, to minimize core body temperature changes. The pupils were dilated with topical 1% cyclogyl-2.5% phenylephrine, and corneas were topically anesthetized with 0.5% proparacaine. For animals analyzed 3-5 hr post induction, anesthesia was re-induced using a ketamine-xylazine mixture of 80mg/1mg/kg. Xylazine’s systemic effects, including bradycardia, hypotension and respiratory depression [23], were reversed using atipamezole (1mg/kg IM). A rat fundus contact lens designed by us and now marketed commercially (Micro-R; Nissel and Cantor, UK) was placed on one eye and coupled using methylcellulose eyedrops. The retina and ONH were imaged using a Haag-Streit model 900 Goldmann slit-lamp biomicroscope (Haag-Streit; Bern, Switzerland). rNAION was induced using an intravenous injection of 1ml/kg of a 2.5mM solution of rose bengal in phosphate-buffered saline (PBS, pH 7.4) administered via tail vein. Thirty seconds post injection, the ONH was illuminated for 6-11 seconds with a clinical 532nM wavelength laser (Iridex Oculight GLx), using a spot size of 500μm diameter, 50mW power. Importantly, because different instruments can have slightly different outputs, or change during time, we measured laser power output at the point of treatment using a laser power meter with a pyroelectric detector optimized for the appropriate wavelength and short duration (1 sec pulse) (Field Mate, Power Max, Coherent, Santa Clara, CA). Laser output varied from 43.5 to 48.1mW at the meter, depending on small variations in angle. Animals were maintained isothermally (38°C) using a circulating heating pad or by deltaphase and were removed from the pad only after recovery from anesthesia to prevent hypothermia that might alter their response to ON ischemic induction. Animals were allowed to recover and used for individual analyses. We previously noted that minimizing the time from rose bengal administration to laser induction is critical for maximizing lesion severity; even as little as a 30-second delay can cause significant changes in the relative lesion severity after induction. However, some interval must exist from dye bolus administration until dye equilibration within the circulation. We used a 30-second ‘mixing time’ from end of dye administration to start of laser induction. rNAION was induced in only one eye of each animal; the other eye was used as an internal (naïve) control. Animals were eliminated from study if there were delays in induction following dye administration.

### Systemic biomarker analysis

Animals were weighed prior to induction, and rose bengal administered according to body weight at 2.5mg/ml/kg. Blood volume in rats is linearly associated with body weight up to 400g [24]. Hematocrit was quantified using capillary centrifugation. Temperature was measured rectally, and heart rate was determined directly using a pediatric stethoscope, with counts obtained at 15 seconds X 4 to yield beats per minute (BPM).

### Local biomarker analysis

#### Fundus color photography

A Nikon D-40 digital camera with slit-lamp adapter was used for color fundus photos. Dye leakage analysis: Both IVFA and ICGA initially were used to evaluate early retina and ONH leakage. We reasoned that the intensity and/or accumulation of fluorescent signal would be an indication of the relative lesion severity, which then could be confirmed by later RGC stereology. IVFA and ICG analysis can be performed using different channels on the Heidelberg SD-OCT instrument, enabling nearly simultaneous IVFA and ICGA pattern comparisons in the same eyes. Animals were induced for 6, 9, 10 or 11 seconds for these experiments.

Five hours post-induction, animals were re-anesthetized and separately intravenously injected with 0.07ml of a 10% solution of sodium fluorescein (fluorescite; Alcon, Geneva, Switzerland) and ICG. ICG concentrations were evaluated from 0.3 to 2% for optimal leakage signal; we ultimately used 0.1ml of a 5mg/ml (0.5%) solution of freshly prepared ICG in sterile dH_2_O for a 250g rat. This corresponds to 2mg/kg body weight. Both injections were independently completed within 30 seconds of each other, as ICG precipitates in saline, and co-administration with fluorescein depletes the ICG signal. We imaged the rodent retina *en face* through the same contact lens used for rNAION induction, coupled to the SD-OCT 25-diopter adapter lens. We initially switched between the two fluorescent-dye detection settings on the Heidelberg instrument. Optimizing individual instrument settings for the analysis was critical, as the Heidelberg instrument is capable of varying laser (excitation) power, detector sensitivity, and signal normalization, all of which will alter final signal output. We used both the 30- and 15-degree field examinations. For IVFA, power was 100 and the detector sensitivity initially was set to 50, without normalization. We used a laser output of 50-100% for both dyes and ICG detector set to 50, without normalization. ONH fluorescence was analyzed at 1, 3, 6 and 12 minutes post-injection. Optimized ICG final settings used 6 min, post-ICG injection with a 15-degree field and 50% power.

We attempted to evaluate further rNAION-associated deep focal ONH leakage using OCT tomography, which generates sequential images, each focused at progressively deeper levels. We used the 5-mm depth tomographic settings available on the current Heidelberg device. The focus point was set at the top of the intraretinal vascular bundle at the top of the optic disk. The diffusible nature of fluorescein made localization difficult. Thus, we assessed only ICG signal in later experiments. Induced eyes were compared against eyes where the animal was treated with Digifab, a known inhibitor of CNS edema (see below).

### Analysis of ONH leakage severity following DigiFab administration

We further examined if ONH leakage can be used as a potential early rNAION biomarker by evaluating the effects of DigiFab, a commercial, clinically approved antibody Fab fragment used to treat digitalis toxicity. DigiFab binds to endogenous ouabain (CTS) with high affinity and can inhibit vascular-induced brain edema [25, 26]. Three additional animals were used, and both eyes of each animal were sequentially induced to enable direct comparison with and without Digifab. Following rNAION (11 seconds) induction of the first eye, Digifab was administered IV at 5.6 mg/kg. This dosage was previously determined to bind endogenous circulating ouabain over a 5-10 hour period following rNAION, with a bound:free ratio of >20:1, enabling binding of ~25naM of endogenous ouabain. Animals received a second dose of DigiFab at 5hr post-induction. Two doses were used to extend the potential effect window because Digifab is rapidly eliminated, with a serum half-life of ~6 hours. The contralateral eye was induced 1 week later with the same conditions, but received vehicle. Analysis was performed using OCT tomography as previously described. ONH edema was measured at 2d post-induction by SD-OCT and RGC loss evaluated by stereology at 30d post-induction (see sections below).

### Analysis of mean ONH diameter, rNAION induction severity and maximum edema

Two days post-induction, animals were re-examined following contact lens application at the slit-lamp and by the Heidelberg device using the IR-OCT display. Using the plano-convex contact lens, both retinal *en face* and cross-sectional imaging can be performed. We used a 15-degree scan through the central retina and ONH, and a 25-slice analysis, with 30 scans averaged/slice (Fig 1D-F).

**Fig 1.**
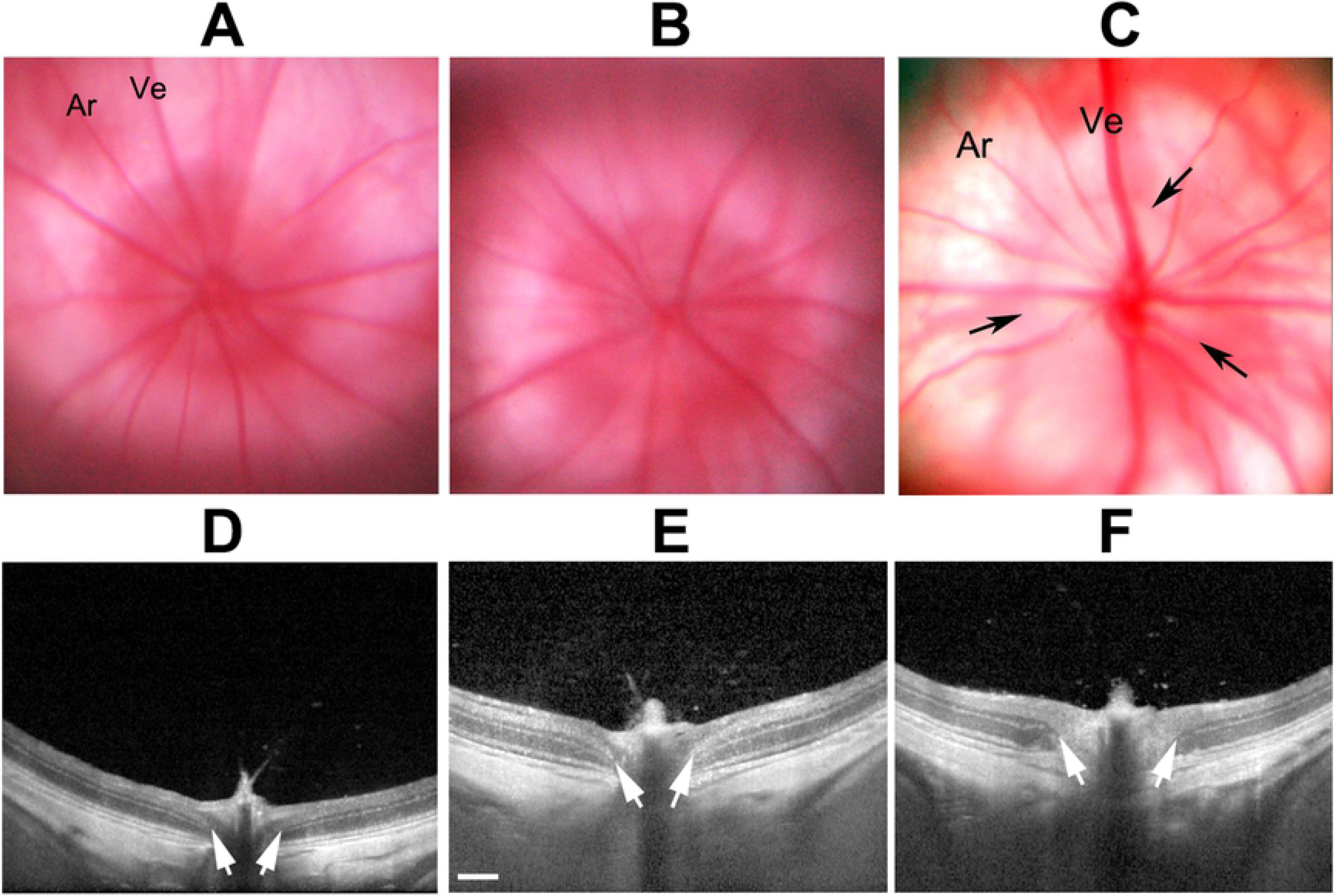
SD-OCT detectable changes following rNAION. **A-C:** Sequential *en face* views of the rodent retina and intraocular portion of the ON in a pre-induced naïve (A), 5hr post (B) and 2d post induction animal (C). **A.** Naïve (uninduced). The ONH has a reddish flush, with the ON border is definable as a slightly lighter color against the retina (Ret). Retinal arteries (ar) and veins (ve) are normal in appearance, and unengorged. **B.**5hr post-induction. The ONH is slightly lighter in appearance, but still flat against the retina. Retinal veins are slightly engorged. **C**. Two days post-induction. The ON is edematous, and pale. The border of the intraocular ON is poorly defined. **D-F:** Sequential cross-section individual slice views of the same nerves shown above, using the Heidelberg instrument and a commercially available rodent fundus contact lens. **D.** Naïve. The intraocular ONH diameter is determined by the INL-INL cross-sectional distance (arrows). **E**. Five hours post-induction. The ONH diameter is slightly enlarged (arrows), and there is engorgement of the vascular tip above the ONH. **F**. Two days post-induction. The ONH diameter is significantly expanded in the intraocular region (arrow), and the nerve fiber bundle has become lighter (edema) and the layers less distinct.

This approach enabled analysis of mean naïve ONH diameter, peripapillary nerve fiber layer thickness, and expansion of the ONH diameter using the instrument-included calipers. The mean ONH diameter was determined by averaging three sequential cross-sections of the ONH with the greatest distance separating both sides of the inner nuclear layer (Fig 1D-1F; arrows).

### Retinal ganglion cell stereology

Thirty days post induction, animals were euthanized, and eyes and optic nerves were isolated and post-fixed in 4% paraformaldehyde (PF)-PBS (pH 7.4). PF-PBS-fixed retinas were isolated, and RGCs immunostained using a primary goat polyclonal anti-Brn3a antibody (Santa Cruz Biotechnology, Dallas, TX), followed by fluorescent labeling with a donkey anti-goat Cy3 secondary antibody (Jackson Immunoresearch, West Grove, PA), flat-mounted and stereologically counted as previously described, using a Nikon Eclipse E800 fluorescent microscope (Nikon, Melville NY) with motorized stage, driven by a stereological imaging package (StereoInvestigator, Ver 10.0; Microbrightfield Bioscience, Williston, VT). Stereological analysis was performed using the Stereo Investigator 10 package, with counts in each eye greater than that required by the Schmitz-Hof equation for statistical validity [27].

## Results

### Systemic parameters in induced animals

Induced animals had a mean rectal temperature of 37.2±0.3° C (range 36.9-37.5° C; n=11). Two animals died after induction (n=2), one of which had a rectal temperature of 35° C and the other, 36.1° C (both considerably below the mean). Mean hematocrit (Hct) was 45.6%, with a range from 43.5 to 51%. The outlier animal with a Hct of 51% had a rectal temperature of 36.9° C and had a minimal (if any) rNAION induction, as measured by SD-OCT mean diameter of 457.3μm and 3.5% RGC loss 30d post-induction. Thus, neither temperature nor Hct consistently correlated with relative induction severity.

### Standard SD-OCT imaging cannot quantify ultimate rNAION lesion severity 5 hours post-induction

We imaged both naïve- and 10-second-induced eyes at both 5hr and 2d postinduction, comparing both the *en face* and cross-sectional appearances of the visualizable ONH using both color fundus photography and SD-OCT. Results are shown in Fig 1.

The naïve eye shows a narrow ONH with a reddish color (Fig 1A, ON). The retinal veins (Ve) are narrow, and the arteries are about ½ the diameter of the veins. Five hours post induction, little color change is apparent in the color fundus photo of the ONH in the *en face* view (compare 1A, naïve, with 1B, 5hr); the amount of ON edema by 5hr is minor, with almost no expansion of the interneural diameter (compare distance between arrows in naïve 1D with those at 5hr post induction, 1E), but with mild blurring of the ONH in cross-section compared with naïve ONH. Two days post induction (Fig 1C, *en face*, and 1F, cross-section) the ONH is visibly edematous (Fig 1C (2d); compare with 1A (naïve). This correlates with expansion of the intraneural diameter (compare 1D [naïve], with 1F (2d post induction). Thus, SD-OCT can detect subtle early (5hr) and later edema-related changes associated with rNAION. We then evaluated the ability of SD-OCT to identify rNAION induction severity.

### rNAION induction reveals an induction threshold effect

The mean ONH diameter of the contralateral (uninduced; naïve) eye was 310±38.2μm (sd), with a range of 254-399μm (n=29 animals). In contrast, the mean ONH diameter of all induced animals induced with the commercially available contact lens (9-11 sec; n=33 animals) was 511±77.2μm, with a range of 359-665μm. We correlated both individual induced-ONH OCT data with subsequent RGC loss data from these eyes, as well as naïve ONH diameter vs RGC loss data from the same animals. Results are seen in Fig 2.

**Fig 2.**
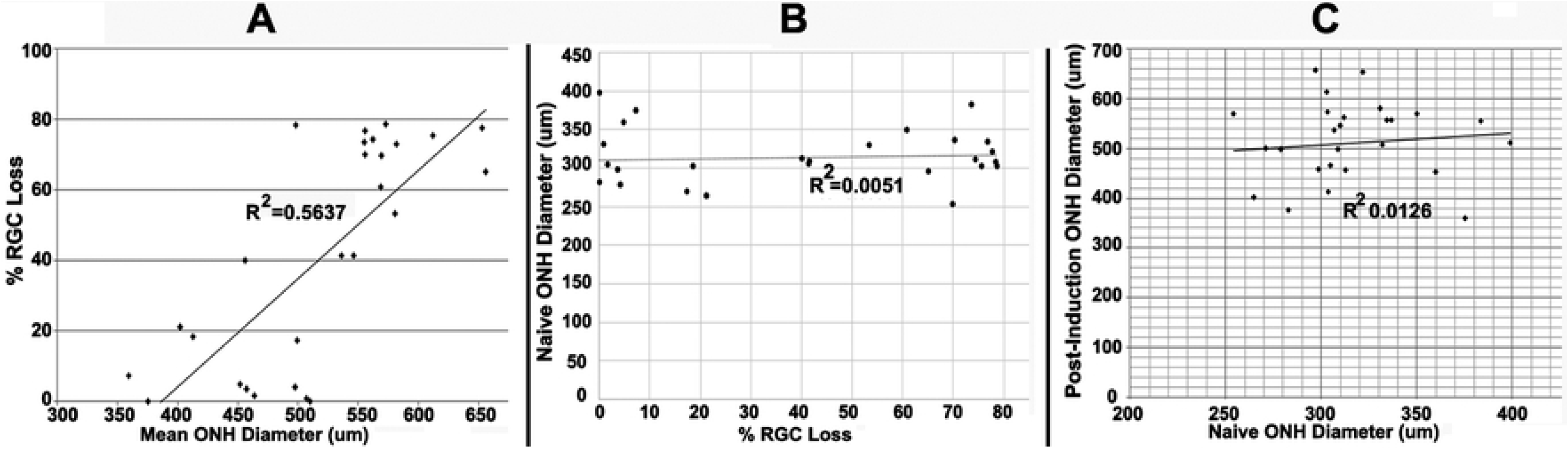
Mean post-induction ONH diameter correlates with RGC loss. A. Correlation of mean post-induction ONH diameter with RGC loss. Post-induction RGC loss is minimal in animals with mean post-ONH diameters <500μm (10/13 animals had RGC loss <20%), whereas relative RGC loss is much greater in induced eyes with mean ONH diameter>500μm; r^2^ is 0.56. **B**. No correlation of pre-induction ONH diameter with subsequent RGC loss;.r^2^ is 0.00. **C**. No correlation of ONH diameter between naïve eye and 2d post-induced eye. The r^r^ value is 0.01.

Analyses were subjected to linear regression, and an r^2^ value derived for each analysis. Comparing individual post-induction ONH edema levels with relative RGC loss reveals a moderate correlation (r^2^=0.5637) of edema severity with RGC loss (Fig 2A). Animals with a mean INL-INL diameter <500μm (189.6μm greater than mean naïve diameter of 310.4μm) had a mean RGC loss of 16.1±5.44% sem (range, 0-78.3%, n=14 animals; 13 animals with RGC loss <25%). In contrast, eyes with a mean INL-INL diameter >500μm had a mean RGC loss of 56.2±5.6% sem (range 0-78.6%; n=19 animals; two animals with RGC loss <25%). This sharp delineation is consistent with a threshold phenomenon associated with a compartment syndrome. In contrast, correlating naïve eye ONH diameter with the percentage of RGC loss in the induced contralateral eye reveals no association between ONH diameter prior to induction and the potential degree of RGC loss following induction (Fig 2B; r^2^=0.0051). Unlike clinical NAION, pre-induction (naïve) ONH diameter does not appear to contribute significantly to the degree of severity of the compartment syndrome generated 2d post induction in the rNAION model (Fig 2C; r^2^=0.0126).

### rNAION lesion severity shows a steep response of RGC loss with laser induction time

RGC stereology using Brn3a(+) staining of retinal flat mounts of naïve eyes revealed a mean count of 1539.004±36.3 (sem) Brn3a(+)RGCs/unit area with a calculated mean total number of 77,101±4748 (sd) RGCs, and a range of 66,800-85,733 (n=53 eyes). Mean inter-eye variation (left-right comparison) was ~ 2.5%. There can be as much as 8% (sd) variability among naïve-eye retinal counts from animals in the same groups.

We compared RGC loss with induction times ranging from 6 to 11 seconds (Fig 3). Six-second induction times yielded no appreciable RGC loss (data not shown). Eleven-second rNAION induction times generated the most consistent ONH lesions, resulting in a mean 71.42 ± 2.53% (sem) RGC loss (Fig 3B; black bar on the far right). An induction time of 10 seconds generated ONH lesions with a mean loss of 40.32±7.79% (sem), and 9 seconds of induction generated a loss of 34.50±17.89 (sem). Thus, reduced induction times are associated with decreased RGC loss. However, there is a steep relationship of average RGC loss response to rNAION induction time.

**Fig 3.**
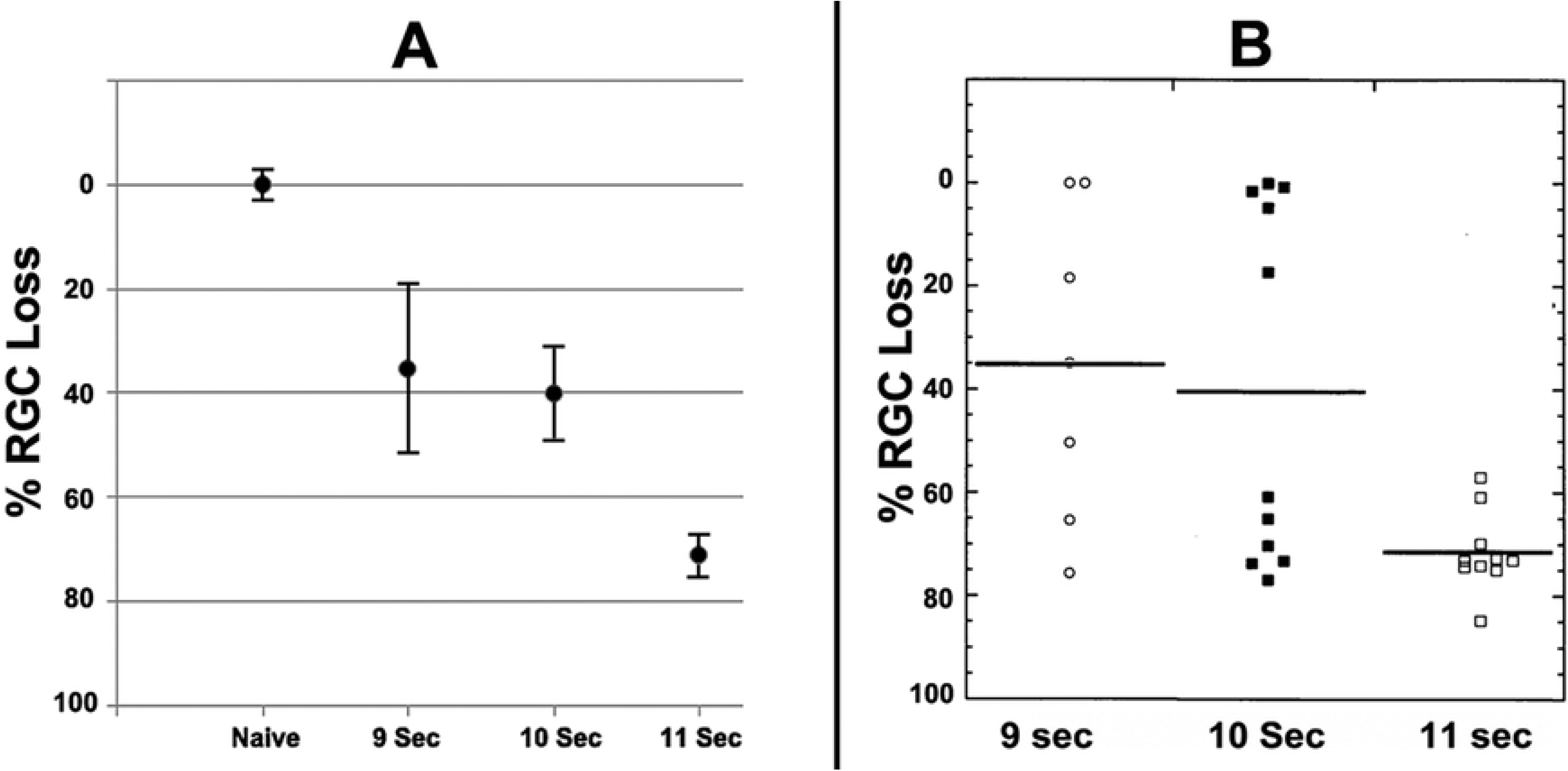
rNAION induction severity in response to laser induction time. **A.** Comparison of RGC counts in animals induced for 9, 10 and 11 seconds. Naïve RGC numbers are given as 100±2.5% (sem). There is a progressive loss of RGCs with increasing laser induction time from 9-11 seconds (Fig 3A). **B.** Scatterplots of individual retinal counts for each eye. There is considerable variability in RGC loss with induction times <11 seconds, and a threshold effect (RGC loss) is clearly discernable in animals treated for 9 or 10 seconds, with ~half of all animals exhibiting significant RGC loss (defined as <20%).

A scatterplot of the data shown in Fig 3B is far more revealing than a simple bar graph. These data reveal that as the relative induction severity (laser induction time) fell below 11 seconds, a significant number of animals in each group failed to show any RGC loss (RGC remaining), with the number of animals without any RGC loss increasing as induction times decreased. For example, nearly all animals induced for 11 seconds had significant RGC loss, (Fig 3B; RGC Loss), but 3/11 (27.3%) animals induced with laser for 10 seconds failed to show any RGC loss (defined as ≤3% difference from the contralateral eye). An even higher 42.8% of animals induced for 9 seconds showed no or only minimal RGC loss, suggesting an rNAION threshold induction effect. We have previously noted that animals induced for 11 seconds or longer occasionally can develop branch or central retinal vein occlusions, resulting in retinal ischemia and associated RGC loss (28), but these complications were not seen in the current study, possibly relating to the use of the commercially available fundus contact lens described above.

### Early Biomarker Analysis using IVFA and ICGA to identify relative post-rNAION lesion severity

Fluorescein is largely unbound to serum proteins and rapidly diffuses through tissue if vascular barriers are disrupted, whereas ICG accumulates near the areas of vascular leakage, enabling identification of different features of vascular decompensation. Following IVFA in naïve (uninduced) eyes, fluorescein signal is constrained within the retinal and choroidal vasculature (Ve and Ar) at 30 seconds post-injection (Fig 4A).

**Fig 4.**
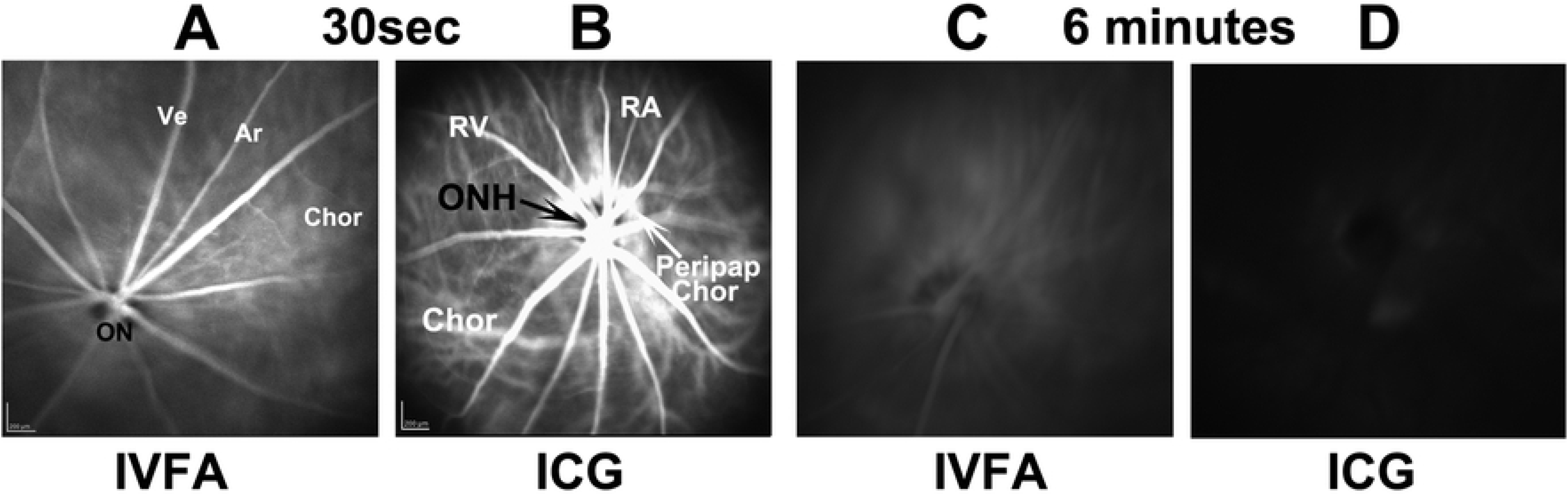
Information obtained through fluorescein and ICG angiography of uninduced (A-D) retinae and ONH. All imaging was done at similar times with the same fundus lens. A and B: IVFA and ICGA imaging, 30 sec post-injection. The IVFA reveals inner retinal vasculature (Ar and Ve). The intraocular optic nerve head (ON) appears as a dark center in the IVFA panel. B. ICG of uninduced eye. The dark ON head (ONH; arrow) is surrounded by a ring of peripapillary choroidal (Peripap Chor) fluorescence surrounding the rat ONH. C and D: IVFA and ICGA photos, respectively, of the same animal, taken 6min post injection. The ON and veins (Ve) are barely visible by IVFA, whereas little if any fluorescence remains in the ICGA channel.

There is poor resolution of the choroidal signal (Fig 4A, Chor). The ONH is dark and remains so in naïve eyes even after several minutes, revealing that fluorescein leakage does not occur in the uninduced ONH. Fluorescein signal is greatly reduced by 6 minutes post-injection (Fig 4C). The ICG signal 30 seconds post IV administration (Fig 4B) reveals a more complex vascular perfusion pattern in both superficial retinal and choroidal vasculature. The choroidal vascular pattern includes additional strong signal in the peripapillary region. Similar to fluorescein, ICG signal rapidly declines after injection; naïve eyes have minimal ONH or retinal signal 6 minutes post injection. Thus, data obtained from ICGA provide more complete vascular information, enabling a more complete analysis.

We evaluated characteristics of rNAION-induced (n=9) eyes for vascular leakage at 5 hours (Fig 5A-F), and ONH edema and fundus appearance at 2 days post induction (Fig 5G-L), comparing IVF and ICG angiography results for each individual animal (shown in pairs: 9-second inductions; A-B; C-D and E-F).

**Fig 5.**
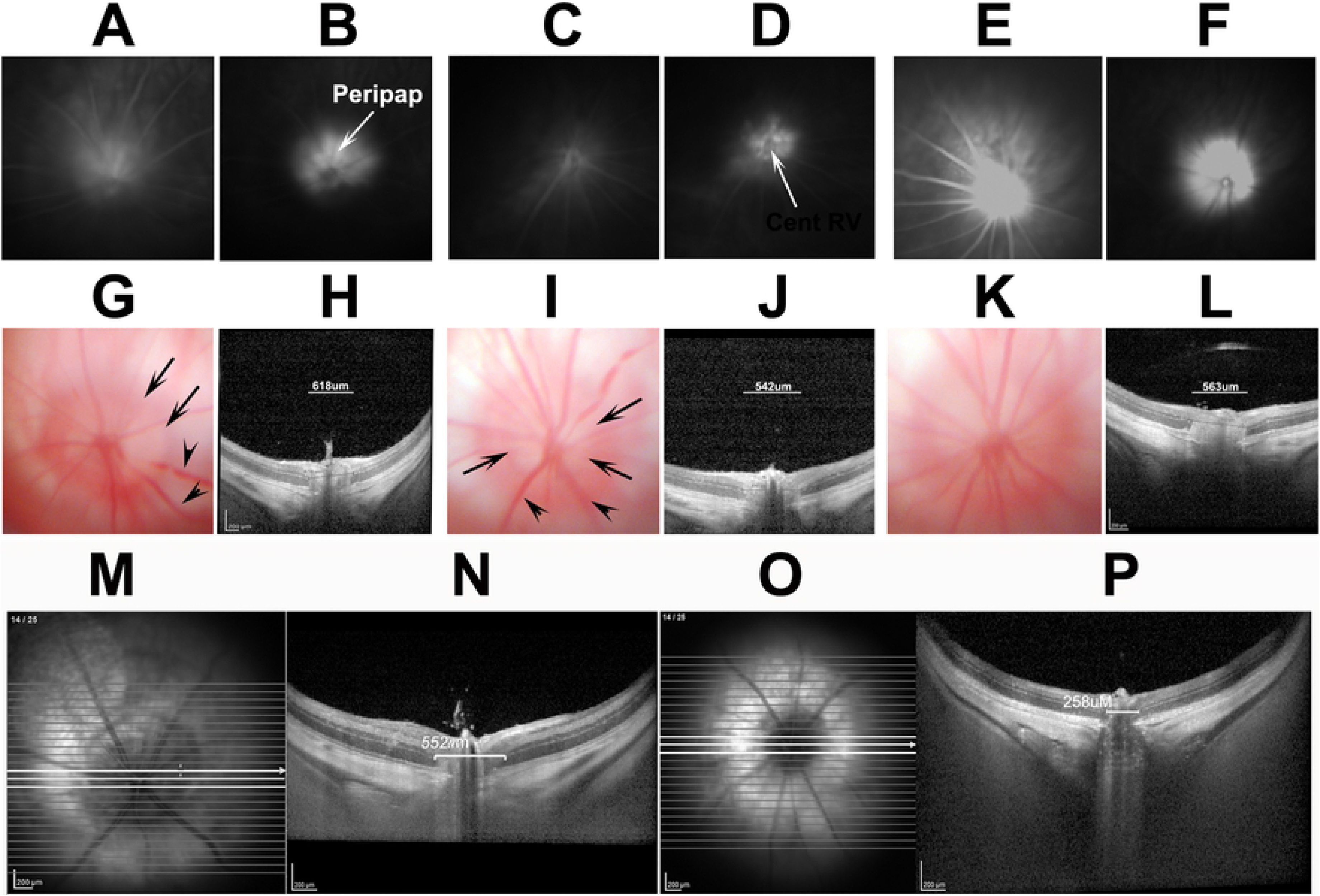
Comparison of characteristics of three similarly induced (9 second) rNAION animals. All animals evaluated at the same time with the same lens and parameters. A-F: paired IVFA (A,C,E) and ICGA (B,D,F), respectively, of three induced eyes 5hr post-induction (ICG 5mg/ml/kg). Diffuse fluorescent signal is present following fluorescein administration above the disk in all three animals, while the ICGA photos reveal localized ICG leakage pattern limited to the ONH, central retinal vascular bundle and the peripapillary RPE. The inner retinal vasculature is apparent in one animal by fluorescein (E; compare with A and C). G-L: Two day post-induction color fundus photo-(G,I,K) and SD-OCT-based cross-sectional image-(H,J,L) pairs taken at the ON centers from the same eyes seen in A-F. There is apparent ON edema in all three eyes (G,I,K) and demonstrable, but varied amounts of expansion of the ON cross-sectional diameter of all three eyes (H,J,L; individual quantification bars seen above the ON). There is also dilation of some retinal veins (arrowheads, G and I) and blurring of the vasculature (see arrows in G and I). There was 46.3%, 60.1% and 28.3% RGC loss at 30 days in these three animals by stereology. M-P: examples of 2d post-induction edema quantification. Following a 150 segmented SD-OCT scan using a fundus contact lens, the INL-INL distance from the three middle cross-sections with the greatest distances are averaged. M and N: imaging from a 9 second eye. O and P: Imaging from the contralateral uninduced eye. Scale bars in M-P: 200μm.

Five hour fluorescence patterns were correlated with SD OCT-based ON edema measures 2d post induction and RGC loss at 30d. This enabled us to determine if relatively early dye leakage correlated with lesion severity and RGC loss, as well as to compare the relative ease of interpretation for the two dyes. All eyes induced for 9 seconds showed ONH leakage at 5 hours, >500μm diameter of ONH edema at 2d, as measured by increase in the mean INL-INL gap (618μm, 542μm, and 563μm respectively) (panels 5H,J,L) and variable but significant RGC loss (28.3, 60.1 and 28.3%), respectively. The 9-second-induced eyes with the lowest fluorescent signal (Fig 5C) yielded more RGC loss (60.1%) than similarly induced eyes with more leakage signal (compare 5A and E; RGC loss 46.3 and 28.3%, respectively), An eye induced for 6 seconds (Fig 6A) with a leakage signal similar to that of an eye induced for 9 seconds had no RGC loss at 30 days (compare Fig 5A, 9 sec, with Fig 6A. 6 sec). Thus, the amount of fluorescein leakage has little correlation with the degree of RGC loss.

**Fig 6.**
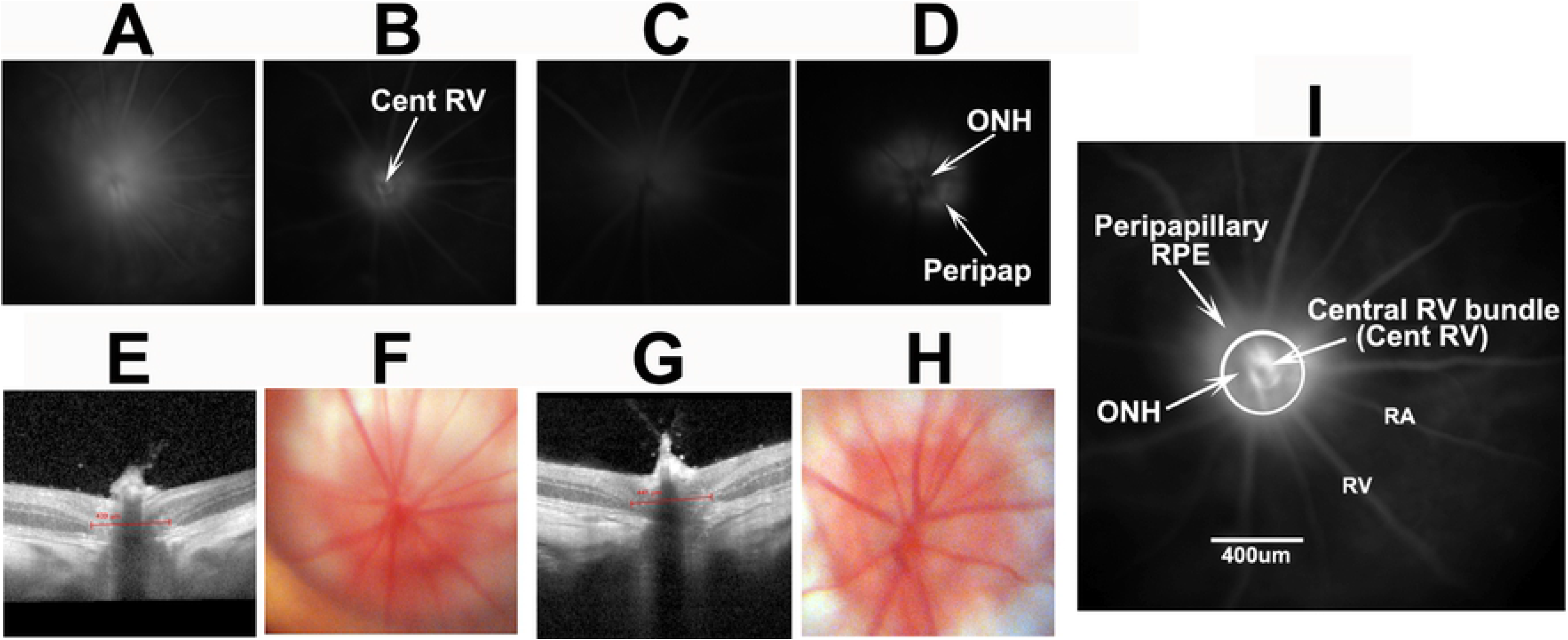
Analysis of two 6-second rNAION-induced animals (no RGC loss) at two days post-induction. A-D: 5hr leakage analysis of fluorescein (A and C) and ICG (B and D). Moderate diffuse fluorescein leakage is seen in one animal (A), and almost none in the other animal (C), while IGC signal at the same time post-injection reveals a more complex leakage pattern, with dye localization around the ONH. E and G: SD-OCT cross-sectional photos. F and H: Fundus color photos. ONH edema is apparent by quantification of the INL-INL distance (439μm and 441μm, respectively, for F and H). There is mild blurring of the ONH margin in the fundus photo F, and none in H. I. Higher magnification of panel 5D, with increased contrast and signal showing ICG leakage. The circle indicates roughly 500um. There is mild signal at the central vascular bundle (site I) in the center of the ONH. The ONH (site II) itself is relatively dark against the surrounding area, revealing reduced leakage relative to the peripapillary RPE (peripapillary sub-RPE) (site III). Scale bar in I: 400μm.

We also evaluated the relative degree of vasculature leakage associated with milder induction (6 seconds) (Fig 6), all of which had no 30d RGC loss. There was less ICG leakage signal at 5 hours (Fig 6B and D), compared with 9-second-induced animals (compare 6B and 6D, 6 seconds with Fig 5B and D, 9 seconds), but vascular leakage at 6 seconds also was highly variable. Six-second-induced eyes also showed less ONH edema at 2 days, with ONH expansion below 500μm (439μ and 441μm, respectively), and minimal visible fundus changes. Fluorescein dye leakage was relatively nonspecific, with poor resolution of the sites of leakage.

A closer evaluation of ICG dye leakage revealed a complex leakage pattern response at 6 minutes post injection in both 9- and 6 second inductions (compare Fig 5B, D, F, and 6B and D).

This is best described by the enlarged ONH-ICG photo in Fig 6I. The strongest leakage signal was generated by peripapillary choroidal vasculature surrounding the ONH under the retinal pigment epithelium (RPE), but there also was ICG leakage from the ONH, compared with its surround (5D; ONH, arrow). Fluorescent signal also was present at the center of the ONH by the central retinal vascular bundle (Fig 6I, Cent RV). Thus, the rNAION induction model generates a complex vascular decompensation response with contributions from the three components: the central vessels emerging from the ONH, the ONH itself, and the choroidal tissue surrounding the ONH, which normally is obscured by the RPE.

In contrast to 9 second inductions, 6-second induced eyes 2d post induction had gross minimal changes in the ONH and surrounding peripapillary retina by slit lamp visualization at 2 days (Fig 6F and H), with minimal ONH edema (defined as <500μm, 439μm and 441μm, respectively; see panels 5E and 5G) and no discernable RGC loss. Naïve (uninduced) eyes had a mean ONH INL-INL gap width of 327.5μm. We also used ICG-coupled ONH tomography, which can be used to image tissue planes at defined depths above and below a particular focal fixation point. Tomography with a 5mm range focused on the anterior posterior pole tissue did not allow better dissection of the leakage at different depths. Thus, ICGA did not enable appreciably better early prediction of ultimate RGC loss than did FA, despite providing additional information about leakage sites.

### ICG-associated ONH tomography and ONH leakage severity following DigiFab administration

DigiFab is a clinically approved antibody that binds to the endogenous Ouabain-like receptor and can inhibit vascular-induced brain edema. Digifab was administered immediately after- and 5hr post-induction. ONH leakage was evaluated at 5hr post-induction via *en face* and ICG-based tomography. Final detector sensitivity was set at 50, with laser power 100% and a 5-mm depth of focus. The three areas identified in Fig 6I were evaluated. Animals were evaluated for ONH edema (INL-INL) at 2d post induction, and RGC counts were generated by stereology. ICG leakage at 5hr and RGC loss (%) at 30d are shown in Fig 7.

**Fig 7.**
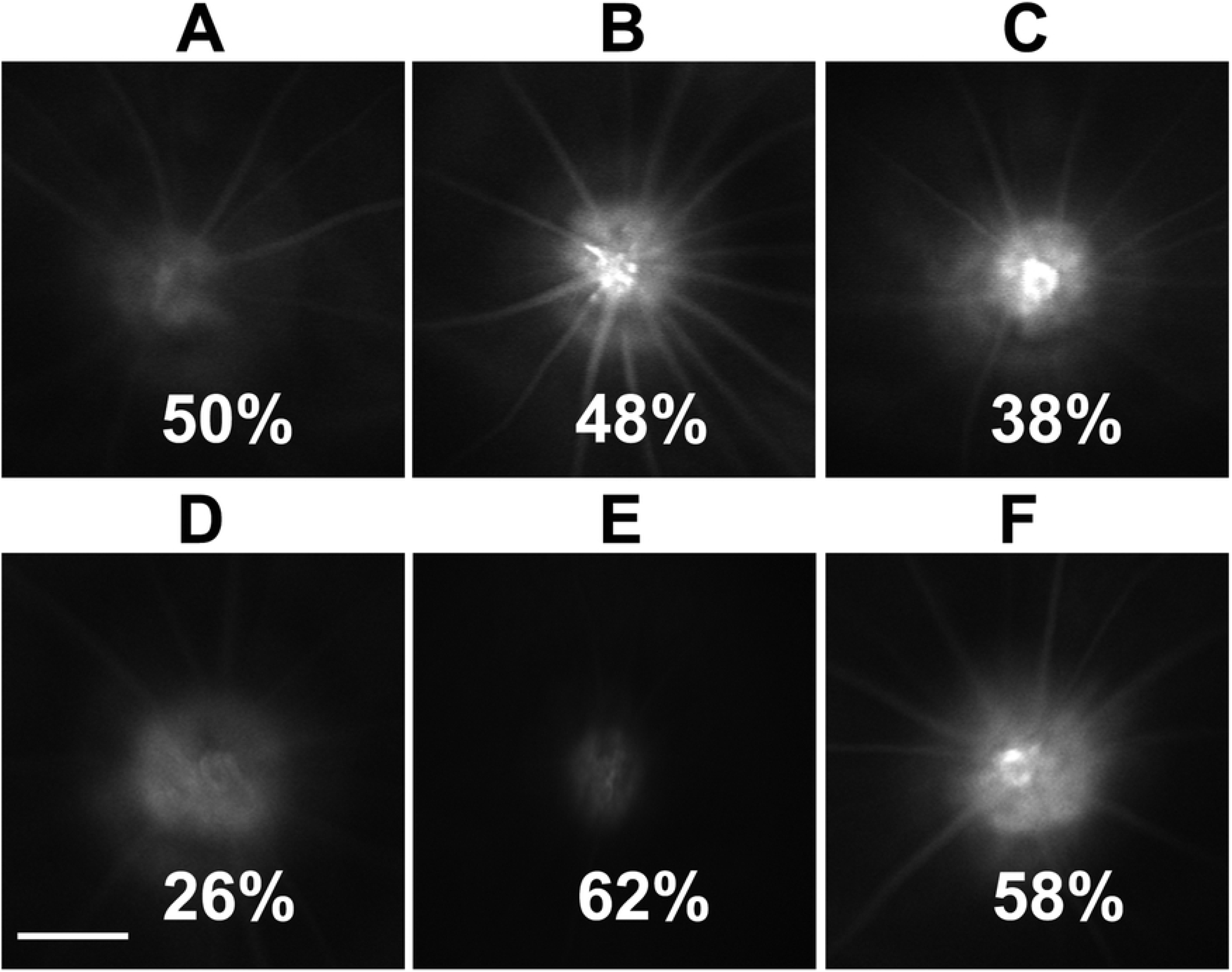
Tomographic ICG-based ONH leakage analysis at 5 hours does not allow RGC loss prediction. Both eyes of each animal were sequentially (1 week apart) rNAION induced for 11 seconds, and ICG (5mg/kg) administered at 5hr post-induction. Tomographic photos were taken at maximal signal point. A-C: Digifab treated. D-F: Vehicle-treated. Each pair of ONHs (A-D, B-E, C-F) are from a single animal.

No consistent pattern of leakage reduction intensity was found, relative to the amount of 30d RGC loss. DigiFab did not demonstrably alter 5hr leakage, 2d post-induction ONH edema, or dramatically change mean RGC loss in Digifab-vs vehicle-treated eyes (47.2±5.4% vs 52.2±16.6% (sd)).

## Discussion

Similar to human NAION, the rNAION model expresses significant variability in its severity and expression in different individuals. This occurs despite maximum consistency of the rNAION induction technique, including optimizing thermal support, using a commercially generated lens, limiting animals to a single sex, normalizing the injected dye volume, maintaining a consistent time from IV dye injection to laser induction, and optimization of laser focus by direct observation. We found no evidence of an association between lesion severity and hematocrit, core temperature, heart rate, or respiratory rate during induction.

The finding that RGC loss in the rNAION model is associated with a threshold expansion of the ONH is consistent with the ‘compartment syndrome’ theory of the pathophysiology of clinical NAION; i.e., there is a threshold level of ONH edema that results in compression of the ONH capillaries in both human NAION and rNAION. However, assuming that both ONHs of each animal are similar in diameter, the rNAION model, unlike human NAION, shows no correlation of preinduction ONH diameter with subsequent ONH edema (r^2^=0.0126), and subsequent RGC loss (r^2^=0.0051), suggesting that, unlike the small cup-to-disc ratio that is the ‘disk at risk’ that predisposes humans to NAION, the rodent model does not possess a homologous association. Our current data suggest that although the rNAION model is similar to human NAION in terms of the development of ONH edema, RGC loss, and other pathophysiological processes such as inflammation [29], the pre-induction diameter of the rodent ONH contributes little if anything to the ultimate development of the pathology. It also should be noted that the animals used in these studies are outbred Sprague Dawley rats. Although this potentially could increase variability in lesion severity, when we performed a small group study using inbred (Wistar) rats, there was no improvement in lesion consistency in terms of measured ONH edema, even when eyes of the same animal were induced sequentially (data not shown). There also were differences in residual RGC numbers between rNAION sequentially induced (1 week apart) eyes of the same animals using the same parameters (data not shown). Thus, there must be other variables that alter the severity of rNAION that have yet to be determined. The accuracy of the SD-OCT device to quantify rodent ONH diameter also is likely less than that for humans, as the Heidelberg device optics and software are designed for human analysis, and the small size of the rodent ONH may result in reduced resolution, despite excellent visualization with the current setup.

Dye leakage studies revealed ONH leakage at 5 hrs post-induction, but these also were not informative with respect to the degree of either lesion severity or ultimate RGC loss. ICG signal enabled considerably more information to be obtained than did IVFA, which largely generated a cloud of dye in front of the ONH. The ICG pattern in uninduced eyes reveals a dense peripapillary choroidal vascularization pattern (Fig 4B), consistent with previous corrosion casting studies demonstrating a major arteriolar supply to the peripapillary choroid that perfuses the ONH [30].

ICG angiography revealed that the vascular-associated leakage that results in the ultimate rNAION lesion is correspondingly complex, with three vascular sites contributing to overall ONH edema: 1) capillaries and larger vessels in the overlying retina; 2) deeper intra-ON capillaries partially obscured by the overlying large retinal veins and arteries (see Fig 4B); and 3) the dense vascular complex within the peripapillary choroid (see Fig 4B). There also was a contribution by the peripapillary RPE, which may be directly affected by laser energy. Disappointingly, the degree of early peripapillary and intraneural dye leakage was not associated with the amount of ultimate ONH edema or RGC loss. This may be due to additional variables yet to be discovered or controlled, such as subtle differences in the angle of laser exposure to the ONH; the insensitivity of current methods to identify the most relevant site of vascular decompensation for progression despite depth sectioning approaches such as OCT and OCTA; or, due to the small size of the rodent ONH, variations in ONH capillary density in individual rodent ONHs that contribute to induction sensitivity. This last possibility seems most likely to us, because mice, with smaller ONHs, have an even greater degree of induction variability [31], whereas there is much less induction variability among eyes in Old World primates, which have considerably larger ONHs and more exuberant capillary beds [32]. Based on this increased morphological size, ICGA leakage analysis may prove useful in identifying regional affected areas in humans who present early in the course of NAION.

Animals in previous studies were evaluated for relative lesion severity at 1-2 days post induction using available methods, including direct visualization and SD-OCT *en face* and cross-sectional analyses. ONH cross-sectional analysis of post-induction ONH edema moderately correlates with ultimate RGC loss (r^2^=0.56). One day post-induction edema values were similar (data not shown). Selecting animals with mean INL-INL expansion >500μm eliminates the vast majority of induced animals with RGC loss <10% (see Figs 1 and 2). Our findings suggest that, like human NAION, rNAION-induced RGC loss likely results from edema causing a compartment syndrome at the ONH and that the minimum required induction parameters exhibit a threshold effect. The ONH diameter-expansion biomarker is likely to prove most applicable in minimizing animal numbers for randomized studies, where treatments begin at least 1-2 days post induction. The majority of studies in this category are neuroregenerative or neuroinflammatory reduction protocols, in which reagents are administered days or even weeks post induction [33]. This approach also may be useful for evaluating early post-induction drug treatment effects such as edema reduction, with sufficient animals, as overall edema is associated with rNAION induction. However, this tool is less likely to be useful in early treatment administration studies, which comprise the majority of current neuroprotective reagents.

Digifab administration had no visible effect on the pattern or severity of early ONH leakage (Fig 7), 2d ONH edema or subsequent RGC loss (Fig 7). This could be due to at least two opposing phenomena: 1) Inhibition of Na pumps by local CTS release could be beneficial by reducing ATP. 2) Inhibiting Na pumps may accelerate the loss of ion gradients, thus enhancing edema formation. These contrasting CTS effects have been previously described in the CNS (34-37) but are beyond the scope of the present study. Additionally, although intravenous administration of DigiFab is clinically the most used route, our results do not exclude the possibility that direct dosing in or near the ischemic area could be beneficial.

In summary, the rNAION model, although useful, must be used with a number of caveats. The model is appropriate in early analyses for identifying edema-reducing therapies, but caution must be taken to include a sufficient number of animals to generate robust data, and induction conditions must be structured to either minimize the number of animals that fail to reach threshold edema levels or include sufficient subjects to overcome this problem. For treatments that can be administered later in the course of the model (ie, neuroregenerative or apoptosis inhibition), ONH edema measurements 1-2d post induction can be used to eliminate non-threshold animals, thus enabling randomization of animals with similar lesion severity into ultimate treatment groups and to reduce the number of study animals that must be carried and processed to final analysis. Other functional approaches, such as OCTA, may prove to be useful in the future for identifying early changes associated with ultimate lesion severity and RGC loss.

## Bibliography

1. Oto S, Yilmaz G, Cakmakci S, Aydin P. Indocyanine green and fluorescein angiography in nonarteritic anterior ischemic optic neuropathy. Retina. 2002;22:187−191.

2. Janaky M, Fulop Z, Palffy A, Benedek K, Benedek G. Electrophysiological findings in patients with nonarteritic anterior ischemic optic neuropathy. Clin Neurophysiol. 2006;117:1158−1166.

3. Kerr NM, Chew SS, Danesh-Meyer HV. Non-arteritic anterior ischaemic optic neuropathy: a review and update. J Clin Neurosci. 2009;16:994−1000.

4. Bernstein SL, Guo Y, Kelman SE, Flower RW, Johnson MA. Functional and cellular responses in a novel rodent model of anterior ischemic optic neuropathy.Invest Ophthalmol Vis Sci.2003;44:4153−4162.

5. Goldenberg-Cohen N, Guo Y, Margolis FL, Miller NR, Cohen Y, Bernstein SL. Oligodendrocyte Dysfunction Following Induction of Experimental Anterior Optic Nerve Ischemia. Invest Ophthalmol Vis Sci. 2005;46: 2716−2725.

6. Chen CS, Johnson MA, Flower RA, Slater BJ, Miller NR, Bernstein SL. A Primate Model of Nonarteritic Anterior Ischemic Optic Neuropathy (pNAION). Invest Ophthalmol Vis Sci. 2008;49:2985−2992.

7. Bernstein SL, Johnson MA, Miller NR. Nonarteritic anterior ischemic optic neuropathy (NAION) and its experimental models. Prog Retin Eye Res. 2011;30:167−187.

8. Tesser RA, Niendorf ER, Levin LA. The morphology of an infarct in nonarteritic anterior ischemic optic neuropathy. Ophthalmology. 2003;110:2031−2035.

9. Pangratz-Fuehrer S, Kaur K, Ousman SS, Steinman L, Liao YJ. Functional rescue of experimental ischemic optic neuropathy with alpha B-crystallin. Eye (Lond). 2011;25:809−817.

10. Huang T-L, Wen Y-T, Chang C-H, Chang S-W, Lin K-H, Tsai R-K. Efficacy of Intravitreal Injections of Triamcinolone Acetonide in a Rodent Model of Nonarteritic Anterior Ischemic Optic NeuropathyTriamcinolone in Rat Ischemic Optic Neuropathy. Invest Ophthalmol Vis Sci. 2016;57:1878−1884.

11. Kapupara K, Huang T-L, Wen Y-T, Huang S-P, Tsai R-K. Optic nerve head width and retinal nerve fiber layer changes are proper indexes for validating the successful induction of experimental anterior ischemic optic neuropathy. Exp Eye Res. 2019;181:105−111.

12. Prieto R, Carceller F, Roda JM, Avendano C. The intraluminal thread model revisited: rat strain differences in local cerebral blood flow. Neurol Res. 2005;27:47−52.

13. Shmonin A, Melnikova E, Galagudza M, Vlasov T. Characteristics of cerebral ischemia in major rat stroke models of middle cerebral artery ligation through craniectomy. Int J Stroke. 2014;9:793−801.

14. Nicholson JD, Puche AC, Guo Y, Weinreich D, Slater BJ, Bernstein SL. PGJ_2_ Provides Prolonged CNS Stroke Protection by Reducing White Matter Edema. PLoS One. 2012; doi:10.1371/Journal.pone.0050021.

15. Ianacone DC, Felberg NT, Federman JL. Tritiated Fluorescein Binding to Normal Human Plasma Proteins. JAMA Ophthalmol. 1980;98:1643−1645.

16. Muci-Mendoza R, Arevalo JF, Ramella M, Fuenmayor-Rivera D, Karam E, Cardenas PL, et al. Optociliary veins in optic nerve sheath meningioma. Indocyanine green videoangiography findings. Ophthalmology. 1999;106:311−318.

17. Yoneya S, Saito T, Komatsu Y, Koyama I, Takahashi K, Duvoll-Young J. Binding properties of indocyanine green in human blood. Invest Ophthalmol Vis Sci. 1998;39:1286−1290.

18. Whittam R. The dependence of the respiration of brain cortex on active cation transport. Biochem J. 1962;82:205−212.

19. Astrup J, Sørensen PM, Sørensen HR. Oxygen and glucose consumption related to Na+-K+ transport in canine brain. Stroke. 1981;12:726−730.

20. Song M, Yu SP. Ionic regulation of cell volume changes and cell death after ischemic stroke. Transl Stroke Res. 2014;5:17−27.

21. Kawamura A, Guo J, Itagaki Y, Bell C, Wang Y, Haupert GT, Jr., et al. On the structure of endogenous ouabain. Proc Natl Acad Sci U S A. 1999;96:6654−6659.

22. Goto A, Yamada K, Ishii M, Yoshioka M, Ishiguro T, Eguchi C, et al. Existence of a polar digitalis-like factor in mammalian hypothalamus. Biochem Biophys Res Commun. 1989;161:953−958.

23. Greene SA, Thurmon JC. Xylazine--a review of its pharmacology and use in veterinary medicine. J Vet Pharmacol Therap. 1988;11:295−313.

24. Lee HB, Blaufox MD. Blood Volume in the Rat. J Nuc Med. 1985;26:72−76.

25. Rap ZM, Schoner W, Czernicki Z, Hildebrandt G, Mueller HW, Hoffmann O. The endogenous ouabain-like sodium pump inhibitor in cold injury-induced brain edema. Acta Neurochir Suppl (Wien). 1994;60:98−100.

26. Wang Y, Fan R, Gu Y, Adair CD. Digoxin immune fab protects endothelial cells from ouabain-induced barrier injury. Am J Reprod Immunol. 1989; 2012;67:66−72.

27. Schmitz C, Hof PR. Design-based stereology in neuroscience. Neuroscience. 2005;130:813−831.

28. Hirabayashi K, Tanaka M, Imai A, Toriyama Y, Iesato Y, Sakurai T, et al. Development of a Novel Model of Central Retinal Vascular Occlusion and the Therapeutic Potential of the Adrenomedullin-Receptor Activity-Modifying Protein 2 System. Am J Pathol. 2019;189:449−466.

29. Salgado C, Vilson F, Miller NR, Bernstein SL. Cellular inflammation in nonarteritic anterior ischemic optic neuropathy and its primate model. Arch Ophthalmol. 2011;129(12):1583−191.

30. Morrison JC, Johnson EC, Cepurna WO, Funk RH. Microvasculature of the rat optic nerve head. Invest Ophthalmol Vis Sci. 1999;40:1702−1709.

31. Bernstein SL, Guo Y. Changes in cholinergic amacrine cells after rodent anterior ischemic optic neuropathy (rAION). Invest Ophthalmol Vis Sci. 2011;52:904−910.

32. Olver JM, Spalton DJ, McCartney AC. Microvascular study of the retrolaminar optic nerve in man: the possible significance in anterior ischaemic optic neuropathy. Eye. 1990;4:7−24.

33. Sozmen EG, Rosenzweig S, Llorente IL, DiTullio DJ, Machnicki M, Vinters HV, et al. Nogo receptor blockade overcomes remyelination failure after white matter stroke and stimulates functional recovery in aged mice. Proc Nat Acad Sci USA. 2016;113:E8453.

34. Jaiswal MK, Dvela M, Lichtstein D, Mallick BN. Endogenous ouabain-like compounds in locus coeruleus modulate rapid eye movement sleep in rats. J sleep Res. 2010;19:183−191.

35. Garcia IJP, Kinoshita PF, de Oliveira Braga I, Parreira GM, Mignaco JA, Scavone C, et al. Ouabain attenuates the oxidative stress induced by lipopolysaccharides in the cerebellum of rats. J Cell Biochem. 2018;119:2156−67.

36. Major S, Petzold GC, Reiffurth C, Windmüller O, Foddis M, Lindauer U, et al. A role of the sodium pump in spreading ischemia in rats. J Cereb Blood Flow Metab. 2017;37:1687−1705.

37. Hodes A, Lifschytz T, Rosen H, Cohen Ben-Ami H, Lichtstein D. Reduction in endogenous cardiac steroids protects the brain from oxidative stress in a mouse model of mania induced by amphetamine. Brain Res Bull. 2018;137:356−62.

